# The generic status of *Anacolosa* (Olacaceae) in Africa with *A. deniseae* a new species to science of Endangered submontane forest liana from Simandou, Republic of Guinea

**DOI:** 10.1101/2022.05.30.493947

**Authors:** Martin Cheek, Denise Molmou, George Gosline, Sekou Magassouba

## Abstract

*Anacolosa deniseae* Cheek (Olacaceae) a submontane gallery forest canopy liana is described as a new species to science and assessed as Endangered using the IUCN 2012 standard due to threats of habitat destruction connected with mining. The roots smell of benzaldehyde when scraped, and the plant reproduces from root suckers. The species is restricted globally to two locations in the Loma-Man Highlands of the Republic of Guinea, all records but one being in the Pic de Fon Fôret Classé of the Simandou Range.

We show that this and the only other continental African species ascribed to the genus *Anacolosa, A. uncifera* of DRC, Gabon & C.A.R., differ in so many architectural, floral and vegetative characters from the remaining species of the genus, which occur from Madagascar to the Western Pacific, including the type *A. frutescens* (S.E. Asia and Indo-China), that they clearly represent a separate genus. The African genus represented by these two species is unique within the Olacaceae (excluding Erythropalaceae) in being a climber (vs. shrubs or trees in *Anacolosa sensu stricto*). Climbing in the two African species is achieved by perennial hook-like structures formed by a combination of five separate traits each of which is unknown elsewhere in the Olacaceae. We formally delimit and describe this new genus, discussing its characteristics, but in the absence of molecular phylogenetic data, refrain from naming it.

## Introduction

In 2006 and 2007 the first author collected specimens of a liana with climbing hook-like petioles from submontane forest in the Simandou and Kourandou Ranges of the Loma-Man Highlands of the Republic of Guinea. The specimens were found by the third author to match a strange species of Olacaceae described from Gabon, Congo and DRC, as *Anacolosa uncifera* Louis & Boutique (Louis & Boutique 1947). In late 2021 the Guinean plant was found again at Simandou in full flower by the second author (Fig. 1), permitting us to test the hypothesis that the Guinean material is a different taxon to that of Central Africa.

**Fig. 1.**
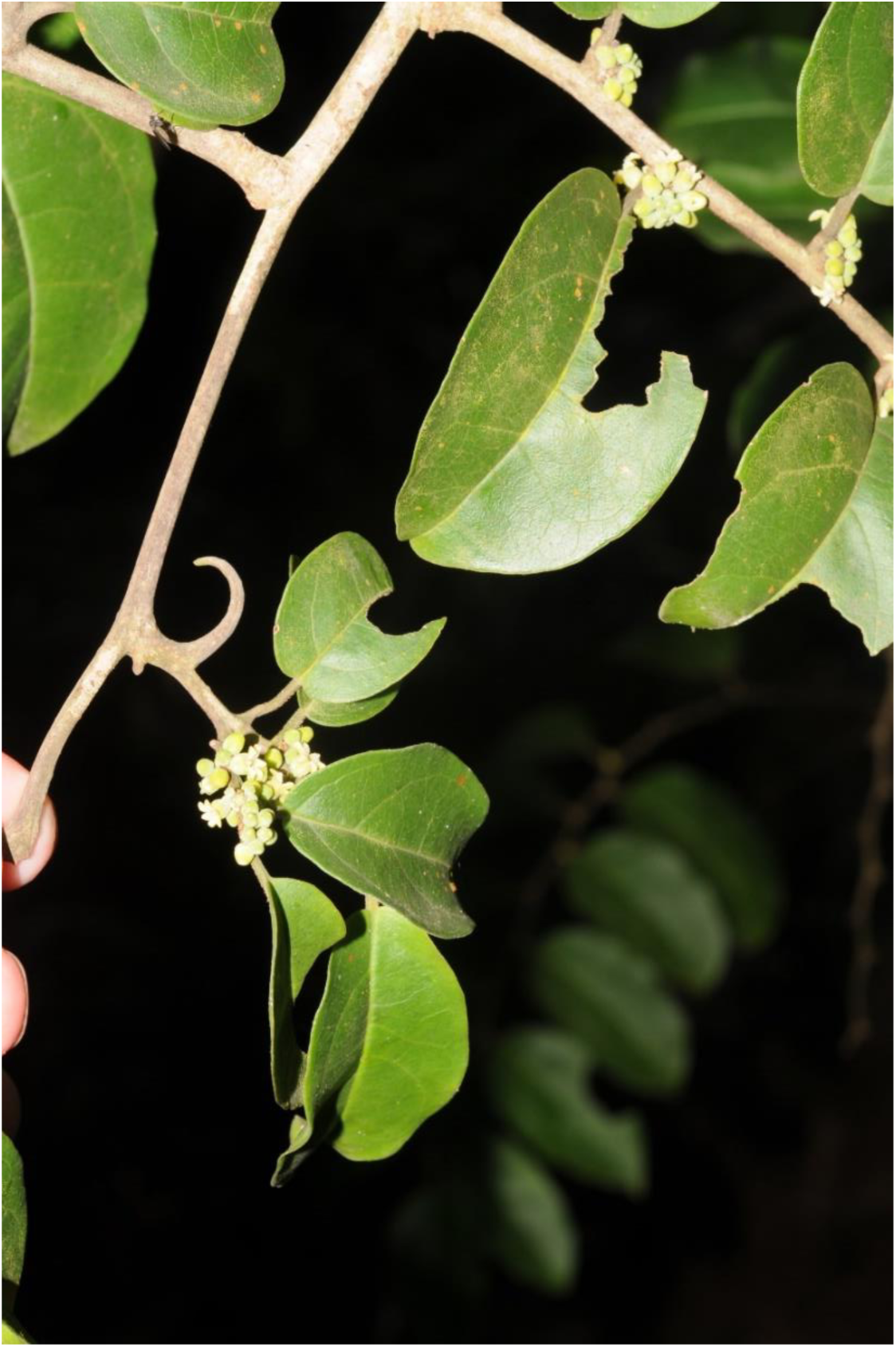
*Anacolosa deniseae* habit, flowering plant, Boyboyba forest, Guinea, Nov. 2021 *Molmou* 1920 (HNG, K). Photo by M. Cheek

Morphological characters separating the Guinean and Congolian material sufficient to support species separation are documented below, and in this paper we name the taxon from Simandou as *Anacolosa deniseae*.

In the course of comparing these two African taxa with each other, we also compared them with the other species placed in *Anacolosa* Blume, which extend from Madagascar to the western Pacific. Although there are similarities between the African and non-African species, there are so many points of difference in habit, foliar structures, indumentum and floral structures that we conclude that the African material represents a new genus to science in the Olacaceae which we describe for the first time but, in the absence of molecular data, refrain from formally naming.

The genus *Anacolosa* was erected for a small tree in Java, *A. frutescens* Blume (Blume 1850). Additional species were added to the genus from S.W. India (Beddome 1864), Indo-China and Myanmar (Kurz 1876; Pierre 1892; Gagnepain 1947; Masters 1875), the Western Pacific (Chrisophersen 1935; Gillespie 1932; Kanehira & Hatusima 1936), New Guinea and neighbouring areas (Sleumer 1980; 1984; Schellenberg 1923), and Madagascar (Cavaco & Kerauden 1963; Baillon 1862). 15 species are accepted (Plants of the World Online, continuously updated).

The non-African *Anacolosa* species are shrubs or trees of lowland evergreen forest, except *Anacolosa pervillei* Baillon, which occurs in semi-deciduous forest in western Madagascar. The centre of diversity is Indo-China, where six species occur, all but one endemic. The western Pacific follows in species diversity, with four species. In contrast Malesia holds only a single widespread species, the type of the genus, apart from New Guinea with two other species (Sleumer 1980).

*Anacolosa* is characterised by six petals, stamens opposite the petals, anthers bearded and placed in the concave lower (proximal) portion of the petals, the upper (distal) petal portions are not papillate (Kuijt & Hansen 2015).

The genus was placed in the Olacaceae tribe Anacoloseae by Engler (1897), together with *Brachynema* Benth., *Strombosiopsis* Engler, *Tetrastylidium* Engler, *Scorodocarpus* Beccari, *Cathedra* Miers, *Strombosia* Blume, and *Worcesterianthus* Merr. (= *Microdesmis* Planch. now Pandaceae) (Sleumer 1935).

Olacaceae *sensu lato* was divided into eight families by Nickrent *et al*. (2010) based mainly on molecular evidence e.g. Malecot & Nickrent (2008). *Anacolosa* was included by Nickrent *et al*. (2010) in the Aptandraceae, together with *Aptandra* Miers, *Cathedra, Chaunochiton* Benth., *Harmandia* Baillon, *Ongokea* Pierre, *Phanerodiscus* Cavaco, and *Hondurodendron* Ulloa *et al*.

However, Kuijt & Hansen (2015), disputed this classification, accepting Aptandraceae, but restricting it to including genera with anthers opening with flaps, and placing *Anacolosa*, which lacks such flaps, into Olacaceae *sensu stricto*, together with *Brachynema, Dulacia* Vell., *Ptychopetalum* Benth., *Strombosiopsis, Olax* L., *Heisteria* Jacq., *Scorodocarpus, Maburea* Maas, *Tetrastylidium, Cathedra, Engomegoma* Breteler, and *Strombosia*.

## Materials & Methods

Specimens were collected using the patrol method as documented in Cheek & Cable (1997). Herbarium material was examined with a Leica Wild M8 dissecting binocular microscope fitted with an eyepiece graticule measuring in units of 0.025 mm at maximum magnification. The drawing was made with the same equipment with a Leica 308700 camera lucida attachment. Specimens, or their high-resolution images, were inspected from the following herbaria: BR, K, P and WAG. Specimens cited have been seen unless indicated “n.v.”. Names of species and authors follow the International Plant Names Index (IPNI continuously updated). Nomenclature follows Turland *et al*. (2018). Technical terms follow Beentje & Cheek (2003). The conservation assessment follows the IUCN (2012) categories and criteria. Herbarium codes follow Index Herbariorum (Thiers continuously updated).

## Results. Taxonomy

Characters separating the Guinean taxon, described below as *Anacolosa deniseae*, from the Central African species *Anacolosa uncifera*, are documented in Table 1, below.

**Table 1.** Characters separating *Anacolosa deniseae* from *Anacolosa uncifera*.

|  | *Anacolosa deniseae* | *Anacolosa uncifera* |
| --- | --- | --- |
| Geography | Republic of Guinea (Guinea Highlands). | Democratic Republic of Congo, Gabon, Republic of Congo, Central African Republic |
| Habitat | Submontane forest 750 – 1300 m alt | Predominantly lowland swamp forest 470 – 700( – 1000) m alt. |
| Leaf-base (flowering stems) | Rounded, cordate or truncate | Cuneate to rounded (never truncate or cordate) |
| Pedicel length | 0.5 – 1 mm | 1 – 1.8( – 2.5) mm |
| Flower bud shape and proportions | Ovoid-globose<br>Length: breadth c. 1:1 | Narrowly conical<br>Length: breadth c. 1.5:1 |
| Flower posture and petals at anthesis | Petals strongly reflexed, splayed horizontally; flower broader than long. | Petals only weakly and slightly reflexed at apex; flower longer than broad |
| Petal lobe indumentum of adaxial surface | Hairs only along midrib at base of lobe | Entire surface densely hairy |
| Anthers | Connective appendage conspicuous | Connective appendage absent |

Louis & Boutique (1947) described *Anacolosa uncifera*, the first continental African species attributed to *Anacolosa* on the basis that the flowers are 6-merous (4-merous or 5-merous in other African genera of Olacaceae), the drupe is enveloped by the accrescent disc and that the calyx is persistent in the fruit. However, we contend that this placement was erroneous and that this species, and the similar Guinean species, described as new to science below, represent a distinct genus of Olacaceae hitherto unknown to science.

The two African species differ from the other species ascribed to *Anacolosa*, including the type species, *A. frutescens*, in numerous characters including some which characterise other genera of Olacaceae (*sensu* Kuijt & Hansen 2015). For example, the corolla of the two African species is united into a tube in the proximal half, as in the S. American monotypic genus *Brachynema*. In the other genera of Olacaceae, including the other species of *Anacolosa*, the petals are either distinct (free) or only basally coherent, not forming a tube.

The two African species also differ from the other species ascribed to *Anacolosa* in that the inflorescence axis resembles that of genus *Strombosiopsis*, in that the flowers are crowded on a swollen, thick inflorescence axis (fig. 2E), the pedicels placed in individual depressions, each subtended by a small bract and two prophyllar bracteoles. Moreover, the anthers in the new species have, like *Strombosiopsis*, a connectival projection (absent in *Anacolosa sensu stricto)*. Although this is not “narrowly triangular” as in *Strombosiopsis* (Kuijt & Hansen 2015), but instead rounded, and resembling an additional, fifth anther theca (Fig. 2 J-L). The African species also have characters otherwise unknown in the Olacaceae, and even in the Santalales. For example, they are lianas, climbing by means of specialised hook-like structures that are developed at nodes that subtend lateral branches. In these petioles the articulation (the abscission zone between petiole and stem), is displaced from the junction with the primary stem (Fig. 2C) towards the distal part of the petiole. The subtending petiole is elongated and reflexes and curves around supports, while its blade is usually caducous (Fig. 2A), but can sometimes persist, although is then often reduced (Hallé 1973).

**Fig. 2.**
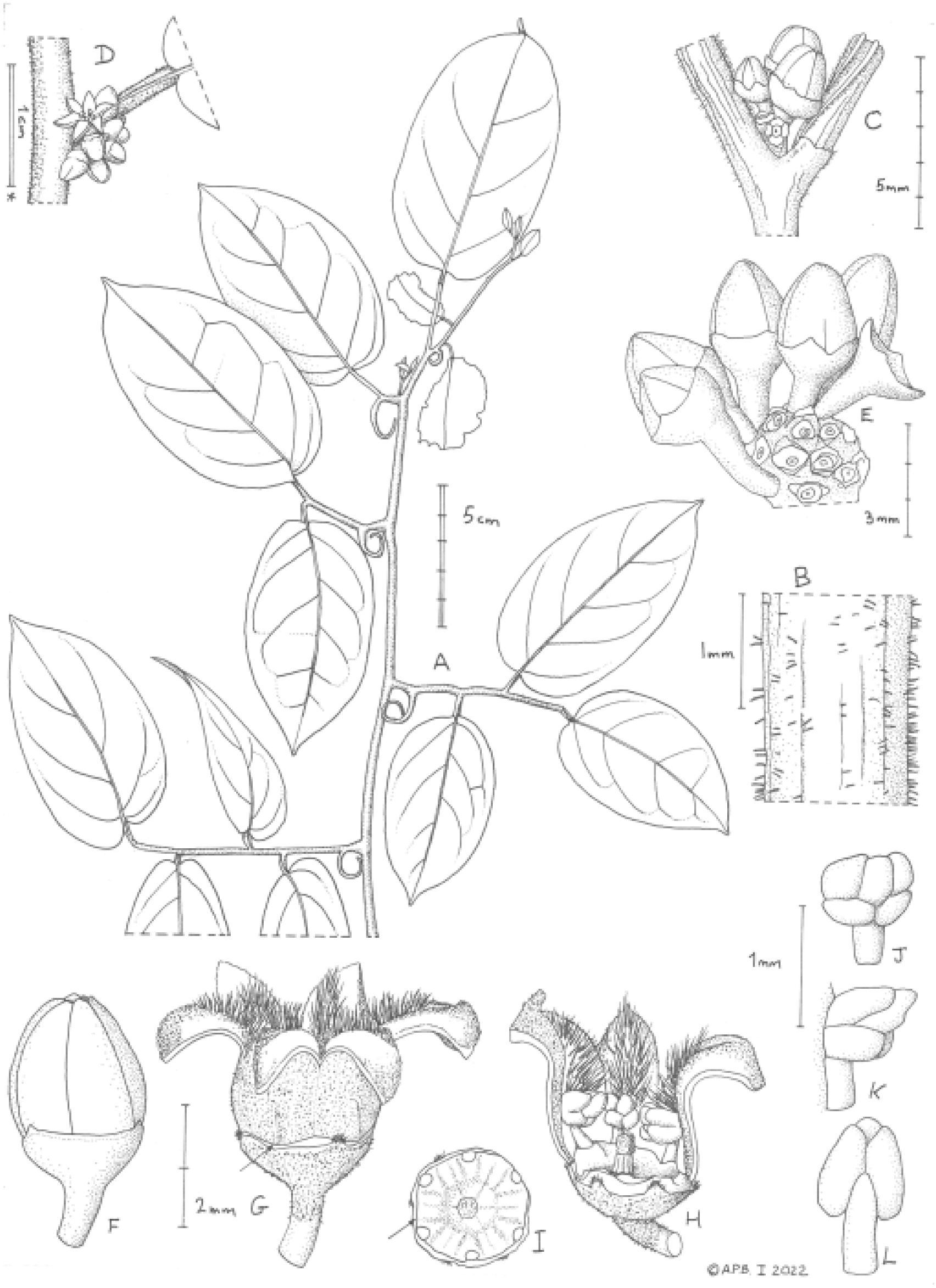
*Anacolosa deniseae* **A** habit; **B** indumentum on stem; **C**. flowering node showing petiole articulation; **D** flowering node with flower at anthesis; **E** inflorescence, globose with sunken sockets; **F** flower bud, side view; **G** flower at anthesis, side view (arrow indicates receptacle-disc); **H** flowers, showing disc, stamens, style-stigma (half of corolla removed); **I** disc, plan view (arrow indicates calyx); **J** stamen (inner face); **K** stamens (side view); **L** stamen (outer face). **A** from *Cheek* 19148A; **B & C** from *Cheek* 13710; **D**-**L** from *Molmou* 1920. Drawn by ANDREW BROWN.

**Fig. 3.**
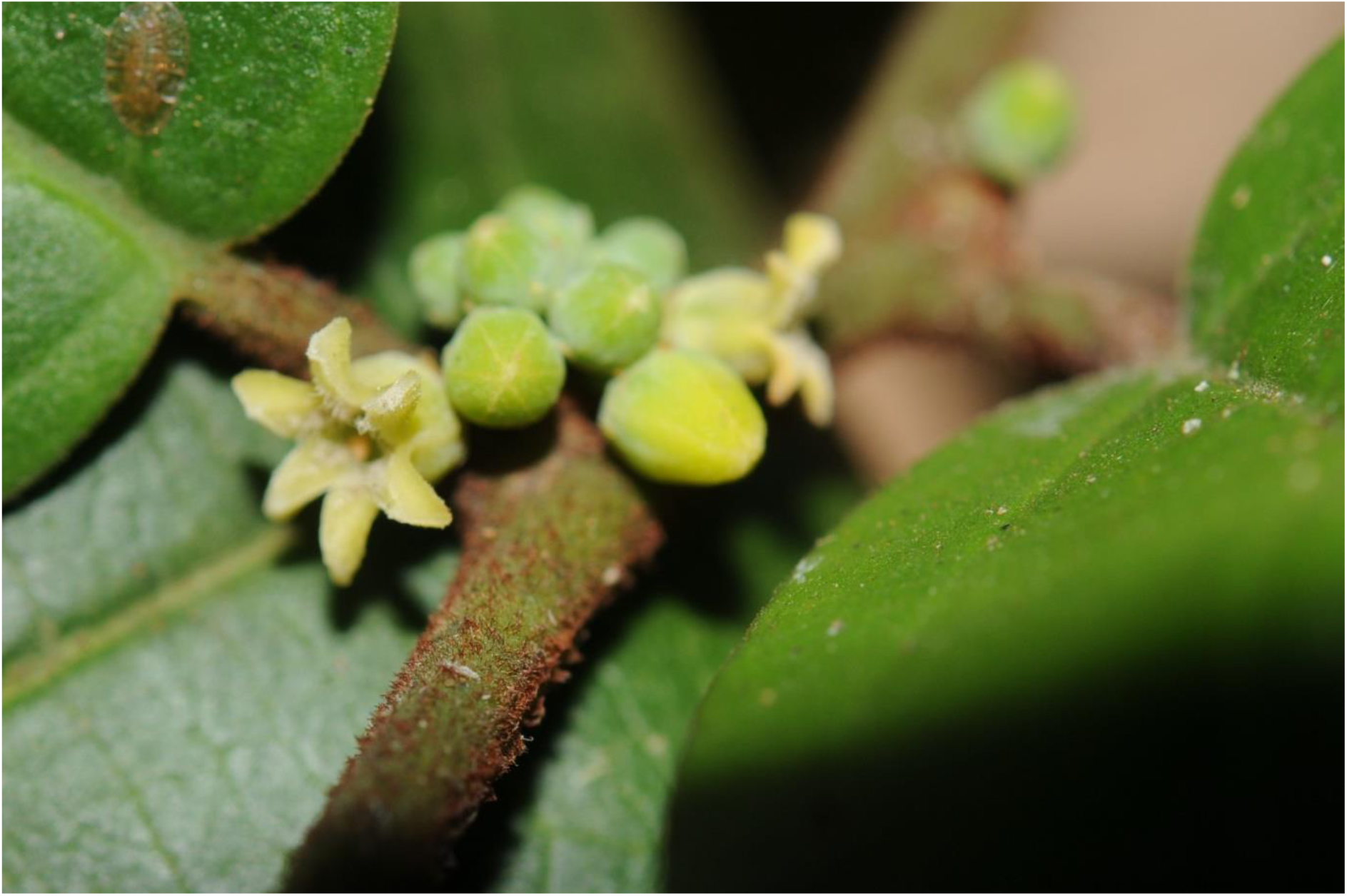
*Anacolosa deniseae* Close-up of flowers at anthesis from *Molmou* 1920. Photo by M. Cheek

**Fig. 4.**
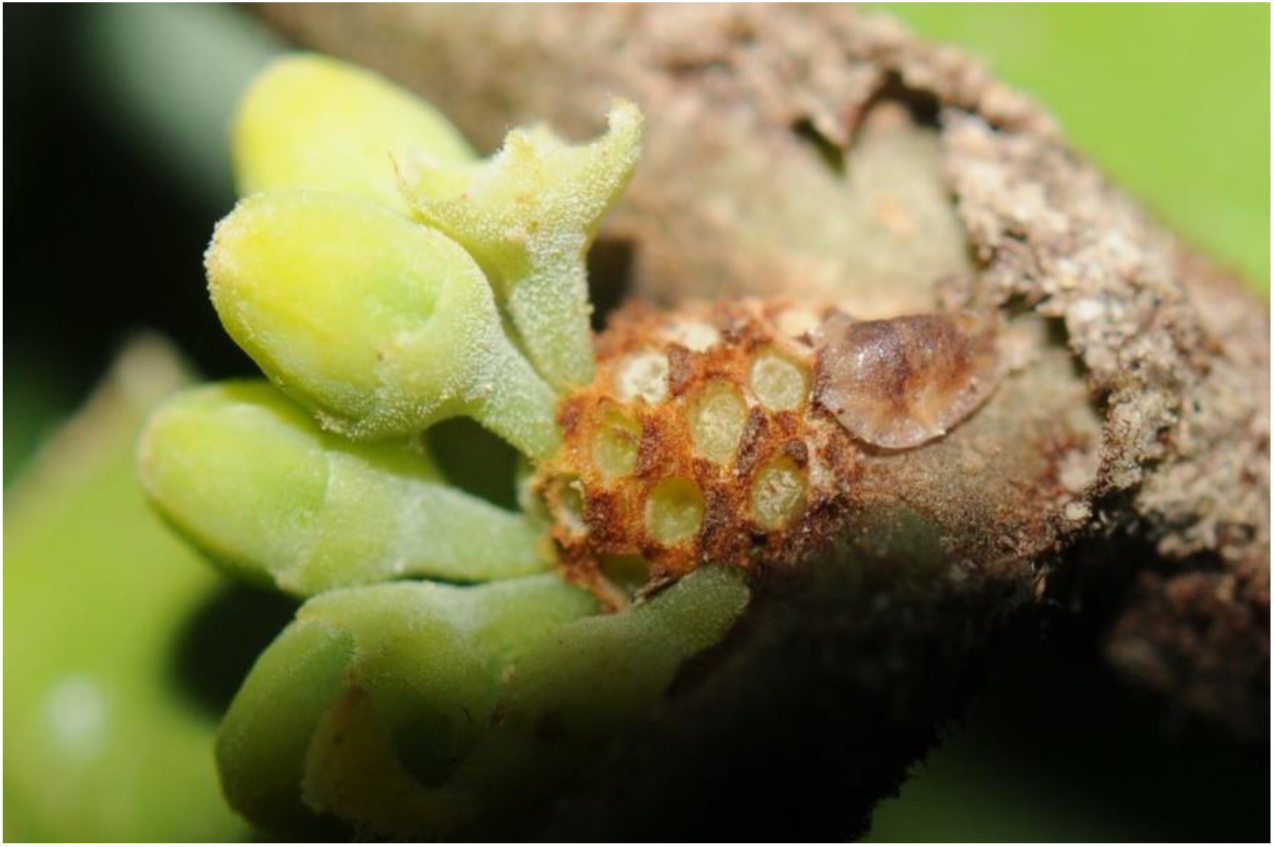
*Anacolosa deniseae* Close-up showing the swollen inflorescence axis with sockets from which originated flowering pedicels. from *Molmou* 1920. Photo by M. Cheek

Remarkably, at the nodes bearing these recurved petioles, the axillary bud and subtending leaf are projected laterally on a 5 – 8 mm stalk (Fig. 2A). Lianas are otherwise unknown in the Olacaceae in the broad sense excepting the monotypic climbing genus *Erythropalum* Blume of S.E. Asia which has tendrils, and which Kuijt & Hansen exclude from Santalales, although Nickrent *et al*. (2010) differ, and include as the monotypic Erythropalaceae. The two *Anacolosa* of Africa are further discordant in the Olacaceae in having multicellular, brown hairs on the stems, petioles and midribs (Fig. 2B), while Santalales generally are glabrous or only papillate, excepting *Coula* Baill. (Coulaceae) and *Octoknema* Pierre (Octoknemaceae).

The African species of *Anacolosa* are also unusual in having leaves distichous, and not spiralled. These, and additional floral features separating the two African species ascribed to *Anacolosa* are outlined in table 2 below, strongly supporting separate generic status.

**Table 2.** Characters separating the continental African species ascribed to *Anacolosa* from those of the Madagascar, Asian and Pacific species.

|  |  |  |
| --- | --- | --- |
|  | African species | Madagascar, Asian and Pacific species |
| Habit & habitat | Forest canopy lianas | Forest understorey shrubs & trees, rarely in woodland |
| Phyllotaxy | Distichous | Spiral |
| Characteristics of principle axis nodes modified for climbing | 1. axillary bud & subtending leaf projected laterally on 5 – 8 mm stalk.<br>2. petiole articulation displaced from base to apex.<br>3. leaf-blade caducous<br>4. petiole reflexed<br>5. petiole twice as long as usual petioles | None of the principal axis nodes are modified for climbing |
| Indumentum of stems, petioles & leaf midribs | Densely red-brown hairy, hairs multicellular | Glabrous or papillate, lacking hairs |
| Inflorescence | Contracted raceme, 6 – 30-flowered, axis thickened, subglobose, flower pedicels placed in individual depressions | Fasciculate or single-flowered, axis neither elongated not thickened, pedicels not in depressions |
| Corolla | Corolla tube subequal to corolla lobes | Corolla tube absent, entirely divided into corolla lobes |
| Staminal filaments | Adnate to corolla tube their entire length | Adnate to corolla lobe at base only |
| Staminal connective | Appendage exceeding anther cells | Appendage not developed |
| Disc | Concave | Convex |
| Style-Stigma | Included in disc | Exserted from disc |
| Stigma | Capitate, not lobed | Lobes 3 or more, acute, short |

## Generic description

### **Genus nov. (**African species of *Anacolosa* Blume)

*Hermaphrodite, evergreen, canopy-flowering lianas, regenerating by root suckers*. Stems often flexuose (zig-zagging). Indumentum dense, red-brown, hairs multicellular, patent, persistent on stems, petioles and midribs of leaves. Phyllotaxy distichous. *Leaves* simple, astipulate, petioles canaliculate, articulate, non-pulvinate, hairs as stems, dimorphic, a) petioles (at nodes of the primary axis subtending axillary stems) projected laterally together with the axillary bud on short stalks several mm long, elongated, reflexing, and articulated at apex, blade caducous; b) other petioles not so elongated, nor reflexing, articulated at base with stem; leaf-blade pinnately nerved, brochidodromous, margin entire, slightly revolute, domatia absent, sparsely hairy near petiole abaxially and on midrib.

*Inflorescences* on axillary stems only, axillary, one per node at multiple successive nodes, racemes condensed, cone-like, 6 – 30-flowered, axis swollen, subglobose to shortly ellipsoid, bracts and bracteole pair triangular, minute, caducous, pedicels inserted in individual shallow pits. *Flower* hermaphrodite. Calyx cupular, shallowly 5-lobed or 5-denticular, inserted at base of ovary-disc, outer surface papillate. Corolla inserted with androecium on rim of the combined ovary-disc, corolla tubular in proximal half, inner surface glossy, glabrous; distal part divided into (5 –)6 valvate, triangular lobes, apex of lobes thickened and triangular in section, adaxial surface (at least part of the midline) long-hairy, otherwise papillate. Stamens 6, opposite the petals, filaments dorsiventrally flattened, glabrous, appressed to the corolla tube their entire length, adnate to base of tube, included in the corolla tube; anthers introrse, inclined inwards, dithecal, the thecae each divided into two superposed, ellipsoid cells arranged around the terminal, distal, connective appendage. Connective appendage ellipsoid, exserted above and slightly larger than the four anther cells, glabrous or sometimes with a few apical hairs. Disc intrastaminal, concave, forming the ovary apex, 5-lobed, lobes broadly rounded, alternating with staminal filament bases, midline of each lobe slightly raised, puberulous. Ovary inferior to corolla + androecium, superior to calyx, 1- or 2- locular, placentation central, apical, ovules 2, pendulous. Style-stigma included within disc, stigma capitate. Fruit a 1-seeded berry, calyx and disc persistent, not conspicuously accrescent, ripening greenish-white, papillate-puberulous, longitudinally shallowly 12-ridged when dry, mesocarp c. 1 mm thick, fleshy; endocarp sclerified. Seed with minute, orthotropic embryo, cotyledons minute, endosperm large. Two species, one throughout the Congo basin extending from DRC into Gabon, CAR, Republic of Congo, the other in Simandou, Guinea Highlands, Republic of Guinea.

#### NOTES

The placement of the new genus represented by the two African species of ‘*Anacolosa*’ remains to be resolved. The differences with *Anacolosa sensu stricto* (the species occurring from Madagascar to the western Pacific, Table 2) are so great that they may not have a sister relationship. The unique climbing mechanism of the African species, with its five component traits unique in Olacaceae, speaks of long evolutionary isolation. It cannot be ruled out that phylogenomic analysis would support family level separation within Santalales for these two species.

Pollen of *Anacolosa uncifera* has been studied by Lobreau-Callen (1980). The grains are triangular-obtuse with straight side and triporate in polar view, 13 – 14 micrometres diameter., rectangular obtuse, flattened in equatorial view, with psilate surface, lacking sculpture.

The Guinean species has been found to reproduce vegetatively, from root suckers. It also produces a strong scent of benzaldehyde from roots and shoots when these are scraped. This information has not been recorded from the Congolian species but should be looked into, it may be that these are additional characteristics of the new genus.

### Key to the species of African ‘*Anacolosa’*

Leaf-blade base cuneate to rounded; abaxial surface when dried orange-brown; pedicel 1 – 1.8(– 2.5) mm long; flower buds narrowly conical. DRC to Gabon… **1. *A. uncifera***

Leaf-blade base rounded, truncate or cordate; abaxial surface drying green to black; pedicel 0.5 – 1 mm long; flower buds ovoid-globose. Guinea **2. *A. deniseae***

**1. *Anacolosa uncifera*** J. Louis & Boutique (1947: 256); Louis & Léonard (1948: 264); Villiers (1973: 104); Hallé (1973: 299).

Lectotype (selected here, see note below): Democratic Republic of Congo (formerly Congo Belge), District Forestier Central, Lumuna, Maniema, forêt, fleurs jaune, Aug. 1932 *Lebrun* 5908 (lecto. BR0000899393; isolecto BR0000899361)

**DISTRIBUTION**. Democratic Republic of Congo, Central African Republic, Gabon, and Republic of Congo

**HABITAT**. Predominantly lowland swamp forest 470 – 700(– 1000) m alt.

**ETYMOLOGY**. Bearing hooks (from the Latin).

**LOCAL NAME & USES**. Kombe Ndjala I Fufow (Turumbu, *Louis* s.n. BR). Uses if any are unknown.

**SELECTED SPECIMENS. Democratic Republic of Congo**, Orientale Prov., Haut-Uele, Asonga Hill, 75 km S of Isiro, outskirts of Ituri Forest, fr. 1 Aug. 2011, *Bujo* 3159 (EPU n.v., K000024628); Maneima Prov. Namoya, Exploration camp to Kibiswa, fl. 9 Aug. 2008, *Q. Luke* 12351 (BR n.v., EA n.v., K, MO n.v.); S Bank of Congo River, st. 8 Nov. 2004, *J*.*P*. & *Q. Luke* 10697Z (BR n.v., EA n.v., K, MO n.v.); Yangole, 20 km W of Yangambi, fl. Feb. 1939, *Louis* 13598 (syntype BR0000899364, BR00008992628); Baringa/S/Maringa (Terr. Befale), foret rivulaire de la Maringa, fl. 25 Oct. 1958 (YBI166807454). **Gabon**. Région de Lastoursville, 1929 – 1931, *Le Testu* 8300 (K, P n.v.)

**CONSERVATION STATUS**. Least Concern. The species has a vast range approximating to the extent of the Congolian forest in DRC, extending over the border into CAR, Gabon and Republic of Congo. The extent of occurrence is calculated as 1,277,662 km^2^, and the area of occupation is estimated as 200 km^2^ using the 4 km^2^ cells preferred by IUCN. Threats to some sites exist e.g. in Gabon, but through most of the range the habitat of this species is not thought to be under pressure and the species is not known to be targeted for exploitation. **NOTES**. In the protologue the authors indicate two specimens as types, one fruiting, *Louis* 13598, the other flowering, *Lebrun* 5908. Of these we here select as lectotype one of the two sheets of *Lebrun* 5908, indicated by barcode above. This is because one of the sheets of this syntype, that chosen as lectotype, bears a floral dissection that is indicated as having been used in the drawing of the protologue, and the metadata agrees with the protologue. The other sheet of this syntype does not, and so becomes the isolectotype.

Since there are some differences in morphology (fruit stipitate vs sessile; connective appendage present vs absent; disc deep vs shallow) between the specimens from Gabon, as depicted in Villiers (1973) and those of DRC as featured in Louis & Boutique (1947), further studies to test the possibility that there may be two taxa rather than one in *A. uncifera* are suggested. In addition, it is notable that while the DRC material described in the protologue derives from inundated and swamp lowland forest, that of Gabon comes from higher, apparently well-drained altitudes. We had insufficient material available from Gabon to investigate this matter during the current study,

**2. *Anacolosa deniseae*** *Cheek* **sp. nov**. Type: Republic of Guinea, Guinea-Forestiere, Simandou Range, Pic de Fon, Fôret Classée, Boyboyba Forest, c. 750 m alt., fl. 20 Nov. 2021, *Molmou* 1920 (holotype HNG; isotypes EA, K000593345, SERG, US) (Fig. 1 – 4)

*Canopy flowering liana*, flowering on stems 5 – 20 m long. *Stems* up to 12 cm diam. at base, older plants with multiple stems arising at ground level, and separate plants arising several metres from the parent from horizontal roots running a few cm below the surface, orangish-white, 1.5 – 2 cm diam; stems woody, cylindrical, bark dark grey-brown, lenticels inconspicuous. Roots and shoots smelling strongly of benzaldehyde when scraped. Leafy stems with principal axis terete, 2.5 – 4 mm diam., internodes (30 –)38 – 50(– 55) mm long; petiolar tendrils (18 –)20 – 25 mm long, nodal protrusions (4 –)6 – 8 mm long; axillary stems up to 38 cm long, 1 – 1.2 mm diam., internodes 9 – 17 mm long, densely, dark brown hairy, hairs multicellular, (0.25 –)0.5 mm long, patent, often with 6 – 7 hairs leaning together, cohering at apex, forming cone-like structures, intermixed with sparse translucent to white papillae 0.05 mm long. *Leaves* drying mid green, concolorous, obovate-elliptic or elliptic 42 – 65(– 82) x (20 –)25 – 36(– 39) mm, acumen triangular (3 –)4(– 6) mm long, base cordate, truncate, or rounded, slightly asymmetric, margin slightly revolute when dried, midrib (and secondary nerves) impressed on upper surface, raised, broad and white on lower surface. *Secondary nerves* 3 – 5 on each side of the midrib, arising at 45 – 70° from midrib, brochidodromous, abruptly angled upwards at junction with secondary nerve below, forming a part looping, part angular infra-marginal nerve 2 – 5 mm from the margin; tertiary and quaternary nerves forming a reticulum with cells isodiametric, c. 1 mm diam. Indumentum of abaxial surface: proximal part as petiole, steadily becoming sparser distally, secondary nerves very sparsely hairy, adaxial surface glabrous. *Petiole* canaliculate (4 –)5 – 8(– 10) mm long, 1.1 mm wide, densely brown hairy as the axillary stems. *Inflorescences* on axillary branches, single in successive nodes, raceme contracted, 10 – 30-flowered, axis swollen, oblate, 1 – 1.5 × 1.5 – 2(– 2.25) mm, bracts caducous, broadly triangular-pentagonal, to 0.5 × 0.75 mm, brown puberulous bracteole pair triangular 0.3 × 0.2 mm, brown puberulous; rhachis brown hairy. *Flower* pedicel 0.5 – 1(– 1.2) mm long, 0.3 mm diam., densely white papillate. *Calyx* broadly cupular, 0.6 – 0.75 × 1.75 (– 2.2) mm, margin with (5 –)6 shallow lobes up to 0.3 × 0.75 mm, or 5 – 6-denticulate, teeth 0.1 × 0.1 mm long, dense brown hairy, indumentum of calyx otherwise densely white papillate; bud broadly ovoid-globose, 2(– 2.5) mm diam. *Receptacle-disc* united at base to calyx, nested within it, cupular, 0.5 – 0.7 × 1.7 – 2 mm, c. 0.5 mm thick, glabrous, outer surface smooth, distal, inner surface of disc with 18 radiating ridges, margin with 6 circular notches, each 0.1 – 0.2 mm diam. (Fig. 2 I & H). *Corolla* inserted on rim of receptacle-disc, white, outer surface moderately densely papillate (50% cover) tube 1 – 1.2 × 2.3 – 2.5 mm, constricted at base and apex; lobes 6, reflexed-patent at anthesis, triangular 1(– 1.5) x 1 – 1.1 mm, midrib raised, lobe base densely long hairy, hairs translucent, moniliform, spreading, 0.4 – 0.5 mm long; remainder of lobe densely papillate, papillae 0.025 – 0.05 mm long, translucent-white. *Stamens* 6, inserted in marginal notches of the disc, opposite to petals and united to corolla tube at base (only), filaments appressed to corolla tube, centripetally flattened, 0.5 × 0.25 – 0.3 mm, glabrous; anthers (inclined centrally Fig. 2K), cordate (Fig. 2 L), 0.5 × 0.75 mm, thecae 2, collateral, each divided into two, subequal, ellipsoid, superposed cells (Fig. 2 J & K); uppermost cells of the two thecae separated by the similarly shaped connective appendage resembling a fifth anther cell (Fig. 2 J-L); anther cells glabrous, or with c. 6 erect hairs 0.35 mm long from the uppermost cells. *Style* subcylindric 0.3 – 0.55 × 0.2 mm, longitudinally ridged, widest at base; stigma capitate, 0.3 × 0.25 – 0.35 mm, minutely pitted (Fig. 2 H). *Fruit* and *seed* unknown.

**RECOGNITION**. Differing from *Anacolosa uncifera* Louis & Boutique in the leaf base cordate or truncate, rarely rounded (vs cuneate to rounded); pedicels 0.5 – 1 mm long (vs 1 – 1.8(– 2.5) mm long); flower bud broadly ovoid to globose (vs narrowly conical). Additional characters in Table 1.

**DISTRIBUTION**. Republic of Guinea: Forestiere Region, Beyla Prefecture, Simandou Range, Pic de Fon Fôret Classée and Kourandou Mts.

**HABITAT**. Canopy-flowering liana of submontane gallery forest, often rooting near streams; 750 – 1300 m alt.

*Anacolosa deniseae* has been collected in flower at the forest-grassland edge (*van der Burgt* 1336). The type material was also collected at the forest edge (*Molmou* 1920). However, the origin of the stems in this last case was found to be downslope, towards a stream, the stems having spread along the canopy to its edge, where (at eye-level) it was more easily detected than high in the forest canopy. Elsewhere in the same forest patch (the Boyboyba forest) more juvenile (non-flowering) plants were seen at the edge of, or near, streams and springs. **ETYMOLOGY**. Named for Denise Molmou, leading Guinea field botanist with the National Herbarium of Guinea and Simfer S.A. who discovered and collected the flowering type specimen material in November 2021. Her commitment and actions for the conservation of threatened species of Guinean plants as part of the Guinea TIPAs project (Couch *et al*. 2019) has been exceptional. She is also commemorated by the endemic, Critically Endangered *Saxicolella deniseae* Cheek (Podostemaceae, Cheek *et al*. in press) for which she collected the type and sole specimen.

**SPECIMENS EXAMINED. Republic of Guinea**. Forestiere Province. **Simandou Range, Pic de Fon Foret Classé**, Fon, st. Aug. 1947 *Schnell* 3307 (PO5327084); ibid. N of Pic de Fon, c. 1.5 km north of Dabatini, fl. 1 Dec. 2008, *van der Burgt* with Pepe Haba and Boubacar Diallo 1336 (HNG, K000460586, P, MO, SERG, WAG); ibid. Moribadou to Canga East 1st. 8 July 2006, *Cheek* with Tchiengue 13326 (HNG, K000437339); ibid above Canga East, st. 17 Nov. 2021, *Cheek* with Tchiam, Molmou, Soropogui 19110 (BR, HNG, K, US); ibid. st. 5 March 2022, *Cheek* with Tchiam, Molmou, Princée, 19213 (HNG, K); ibid.

Oueleba, Boyboya Forest, st. 20 Nov. 2021, *Cheek et al*. 19148 (HNG, K); ibid., fl. 20 Nov. 2021, *Molmou* 1920 (holotype HNG, isotypes K, SERG, US); gallery submontane forest, Oueleba, eastern side, 14-20 Dec. 2021 *Molmou* with Thiam & Soropogui, sight record; Oueleba, northern side, Siatoro forest, Dec. 2021 *Molmou* with Thiam, sight record; **Kourandou Mts**, Sinko to Seberendou path, fl. buds 17 Nov. 2007, *Cheek* with Seydou Cissé, P. Haba and Condé 13710 (HNG, K000615587).

**CONSERVATION STATUS**. *Anacolosa deniseae* is known from only two locations, one at the Kouroundou Mts (*Cheek* 13710, HNG, K) with a single site and individual recorded, where it grows close to a footpath in a farming area near a major town and is at risk of habitat clearance for charcoal production. At the second location, the Pic de Fon Fôret Classée,

Simandou Range, six sites are recorded all with a single individual (including offset suckers, see Notes below) except one, the Boyboyba Forest, with five individuals recorded including the type collection. This location is set to host a major open pit iron ore mine. While the pit is expected not directly to impact sites for this species, there is a risk that despite best efforts to protect threatened species, nevertheless there will be negative impacts linked with the activities and infrastructure associated with the expected ore extraction, such as dumping of waste rock or alteration of hydrology. Extent of occurrence is calculated as 407 km^2^, and area of occupation as 28 km^2^ using the IUCN required 4 km^2^ cell-size. Only three mature (flowering-sized, reaching canopy) individuals are known, accordingly we assess *Anacolosa deniseae* as Endangered EN B1+B2a,b(iii). It is to be hoped that this species will be searched for and found at other locations which would allow a lower extinction risk assessment than that made here. However, since this species is so distinctive and easily recognised even at a juvenile stage it is strange that if it has a wider range, that it has not been detected elsewhere before now.

**NOTES**. *Anacolosa deniseae* is immediately recognisable in Guinea in the field, even as juvenile plants only 1 – 4 m tall, due to the conspicuous and unusual climbing stems (Fig. 1). Seedlings less than c. 60 cm tall lack climbing hooks.

Two seedlings less than 1 m tall, separated by c. 100 m, and two distantly separated larger juvenile plants 1 – 4 m tall were found in the Boyboyba forest of the Simandou range in late 2021, suggesting that natural regeneration from seed is occurring. The means of seed disperal is unknown. We speculate that fruit might be produced as the wet season begins in May.

Searches in March for fruit on the plant found in flower in November did not show even immature fruit suggesting that fruits may not be produced every year.

Researching this species in the field at two sites in March 2022, we found that full-sized plants at both sites had several (3 – 7) satellite plantlets arising from roots that we traced from the full-size parent. Identifying and tracing the roots of this species is facilitated by their strong scent of benzaldehyde, unusual in the plant world. Vegetative apomixis such as we observed is not often recorded in Olacaceae in Africa. The juveniles were all within 4 m of the parent and mostly about 1 – 2 m tall.

*Anacolosa deniseae* at first sight is closely similar to *A*.*uncifera*, differing in the features detailed in Table 1, supporting the large geographical disjunction of 2600 km between these two species. The absence from the Cross-Sanaga Interval, in western Cameroon and SE Nigeria, which has the highest species and generic diversity for plants per degree square in tropical Africa (Cheek *et al*. 2001, Barthlott *et al*. 1996, Dagallier *et al*. 2020) is difficult to explain since many, if not most of the recently discovered Guinea Highland endemics with disjunctions to the east occur in the highlands of the Cross-Sanaga Interval (e.g. in the genera *Brachystephanus, Isoglossa, Talbotiella, Ternstroemia*, respectively Darbyshire *et al*. 2012; Champluvier & Darbyshire 2009; van der Burgt *et al*. 2018; Cheek *et al*. 2019a). This is the first case that we are aware of a far wider disjunction in this context.

## Discussion

### Threatened plant species of the Simandou range

*Anacolosa deniseae* is the latest in a line of species new to science discovered from the Simandou Range, all of which are threatened with extinction and constitute the highest priorities for plant conservation within that range. The species are either from the submontane bowal grassland on ferralitic (iron-rich) substrate (*Xysmalobium samoritourei* Goyder (Apocynaceae, Goyder *et al*. 2009), *Coleus ferricola* Phillipson, O.Hooper & A.J. Paton (Lamiaceae, Phillipson *et al*. 2019), *Polystachya orophila* Stévart & E.Bidault (Orchidaceae, Bidault *et al*. 2016) or from adjacent submontane forest: *Brachystephanus oreacanthus* Champl. (Acanthaceae, Champluvier & Darbyshire 2009), *Isoglossa dispersa* I. Darbysh. (Acanthaceae, Darbyshire *et al*. 2012), *Gymnosiphon samoritoureanus* Cheek (Burmanniaceae, Cheek & van der Burgt 2010), *Allophylus samoritourei* Cheek (Sapindaceae, Cheek & Haba 2016a) and *Psychotria samoritourei* Cheek (Rubiaceae, Cheek & Williams 2016), or restricted to the interface (or transition) between these two habitats: *Hibiscus fabiana* Cheek (Malvaceae, Cheek *et al*. 2020a), while one occurs in mainly lowland woodland: *Striga magnibracteata* Eb. Fischer & I. Darbysh. (Orobanchaceae, Fischer at al. 2011).

Most of these species have since been found in one or more other locations outside the Simandou Range, but other discoveries, (*Eriosema triformum* Burgt (Leguminosae, van der Burgt *et al*. 2012) and *Keetia futa* Cheek (Rubiaceae, Cheek *et al*. 2018a)), remain globally restricted to the Simandou Range on current evidence and are Critically Endangered. The last species formerly occurred at two other sites far from Simandou where it has been lost. *Anacolosa deniseae* will also become restricted to Simandou and need re-assessment, likely as Critically Endangered, if it is lost at its single, unprotected site outside of Simandou near Sinko.

### New plant discoveries elsewhere in the Republic of Guinea

The Simandou Range is not exceptional in Guinea for conservation importance and recent discovery of new, range-restricted species. The Kounounkan area of the Fouta Djalon range, with habitats based on table mountains of Ordovician sandstone is far more exceptional in terms of historically known endemic taxa, such as the monotypic genus *Caillella* Jacq.-Fél. (Melastomataceae, Veranso-Libalah *et al*. 2021), *Mesanthemum benna* Jacq.-Fél. (Eriocaulaceae, Phillips *et al*. 2018), *Impatiens benna* Jacq.-Fél. and *Rhytachne perfecta* Jacq.-Fél. (Gramineae, both Couch *et al*. 2019). Surveys for the Guinea Tropical Important Plant areas programme have recently resulted in discovery of the following range-restricted species, all also threatened: *Keetia susu* Cheek (Rubiaceae, Cheek *et al*. 2018a), *Gladiolus mariae* Burgt (Iridaceae, van der Burgt *et al*. 2019), *Ternstroemia guineensis* Cheek (Ternstroemiaceae, Cheek *et al*. 2019a), *Trichanthecium tenerium* Xanthos (Gramineae, Xanthos *et al*. 2020), *Ctenium benna* Xanthos (Gramineae, Xanthos *et al*. 2021) and even the Kounounkan endemic new genus *Benna alternifolia* Burgt (Melastomataceae, van der Burgt *et al*. 2022), with additional endemic range-restricted species to be described of *Virectaria* (Rubiaceae) and *Hibiscus* (Malvaceae).

Elsewhere in Guinea range-restricted new species to science have been found in numerous provinces and habitats, from surviving fragments of lowland forest, to cliffs, flate bare sandstone, and waterfalls and rapids: *Eriocaulon cryptocephalum* S.M.Phillips & Mesterházy (Eriocaulaceae, Phillips & Mesterházy 2015), *Napoleona alata* Jongkind (Lecythidaceae, Prance & Jongkind 2015), *Inversodicraea pepehabai* Cheek (Podostemaceae, Cheek & Haba 2016a), *Karima* Cheek (Euphorbiaceae, Cheek *et al*. 2016), *Kindia gangan* Cheek (Rubiaceae, Cheek *et al*. 2018b), *Talbotiella cheekii* Burgt (Leguminosae, van der Burgt *et al*. 2018), *Inversodicraea koukoutamba* and *I. tassing* (Podostemaceae, Cheek *et al*. 2019b) and *Vepris occidentalis* Cheek & Onana (Rutaceae, Cheek *et al*. 2019c).

## Conclusions

About 2000 new flowering plant species are described each year (Cheek *et al*. 2020b), adding to the estimated 369,000 (but number debated) already known to science (Nic Lughadha *et al*. 2016; 2017). Widespread species tend to have already been discovered, although there are exceptions, such as *Vepris occidentalis* (cited above) that occurs from Guinea to Ghana. More usually, newly discovered species are those that are range-restricted and so are much more likely to be threatened, such as *Anacolosa deniseae* published here. Until new species are formally named and known to science, it is much more difficult to assess them for their IUCN conservation status and so the possibility of protecting them is reduced (Cheek *et al*. 2020b). The majority of plant species still lack such assessments (Nic Lughadha *et al*. 2020). Documented extinctions of plant species are increasing (Humphreys *et al*. 2019) and recent estimates suggest that as many as two fifths of the world’s plant species are now threatened with extinction (Nic Lughadha *et al*. 2020). This makes it imperative to discover and publish such species so that they can assessed, and, if merited, conservation actions taken to avoid the risk of becoming, like Guinea’s *Inversodicraea pygmaea* G.Taylor, globally extinct (Cheek 2018, Cheek & Magassouba 2018). Designating and implementing Important Plant Areas (Darbyshire *et al*. 2017; continuously updated) is key to *in situ* conservation of plant species. For this reason the Important Plant Areas (TIPAs) of Guinea have been recently designated (Couch *et al*. 2019) and accepted by the Government of Guinea (Col. Seyba, head of Oguipar (protected areas) pers. comm. 2019). Fortunately, *Anacolosa deniseae* occurs within the newly designated Southern Simandou TIPA (TIPAs Guinea-Conakry (2016 – 2019); Couch *et al*. (2019).

The authors declare no conflict of interest.

## Acknowledgements

This paper was completed as part of the Guinea Important Plant Areas programme managed by Charlotte Couch, currently supported by the Franklinia Foundation and the Critical Ecosystem Partnership Fund (CEPF).

Most of the specimens cited were collected with the support of Simfer S.A. and those collected in 2021 and 2022 under the framework of collaboration between RBG, Kew and Sylvatrop Consulting. We especially thank former Simfer Environment staff John Merry, Leon Payne, Thomas Williams, and current staff Mohamed Tahlaoui, M. Soumaoro and M. Diaby. At Sylvatrop, we thank especially Dr Eric Muller.

The Bentham Moxon Trust through the efforts of Dr Isabel Larridon, supported Denise Molmou’s visit to RBG Kew in April-June 2022 during which time it was possible to complete the paper. The authors thank their colleagues at Université de Gamal Abdel Nasser-Herbier National de Guinée (MINRESI), for support. Two anonymous reviewers are thanked for constructive comments on an earlier draft of the paper.

